# Quantifying the contribution of subject and group factors in brain activation

**DOI:** 10.1101/2022.08.01.502338

**Authors:** Johan Nakuci, Jiwon Yeon, Kai Xue, Ji-Hyun Kim, Sung-Phil Kim, Dobromir Rahnev

**Affiliations:** School of Psychology, Georgia Institute of Technology, Atlanta, Georgia, 30332, USA; Department of Psychology, Stanford University, Stanford, California, 94305, USA. California, 94305, USA; Department of Biomedical Engineering, Ulsan National Institute of Science and Technology, Ulsan, South Korea

**Keywords:** individual differences, fMRI, perceptual decision-making, brain-behavior relation

## Abstract

Research in neuroscience often assumes universal neural mechanisms, but increasing evidence points towards sizeable individual differences in brain activations. What remains unclear is the extent of the idiosyncrasy and whether different types of analyses are associated with different levels of idiosyncrasy. Here we develop a new method for addressing these questions. The method consists of computing the within-subject reliability and subject-to-group similarity of brain activations and submitting these values to a computational model that quantifies the relative strength of group- and subject-level factors. We apply this method to a perceptual decision-making task (N=50) and find that activations related to task, reaction time (RT), and confidence are influenced equally strongly by group- and subject-level factors. Both group- and subject-level factors are dwarfed by a noise factor, though higher levels of smoothing increases their contributions relative to noise. Overall, our method allows for the quantification of group- and subject-level factors of brain activations and thus provides a more detailed understanding of the idiosyncrasy levels in brain activations.

## Introduction

Human behavior is idiosyncratic: what elicits a certain behavior in one person is often very different from what elicits that same behavior in another (Eilam 2015; Forkosh et al. 2019). Similarly, increasing amount of evidence points towards the existence of substantial idiosyncrasy in brain activations, such that the same task can elicit different patterns of activity in different subjects (Seghier et al. 2008; Miller et al. 2009, 2012). Yet, it remains unclear how to precisely quantify the strength of the observed idiosyncrasy, as well as whether different types of analyses are associated with different levels of idiosyncrasy.

To address these questions, here we develop a method to determine the contribution of group- and subject-level factors to observed activations in functional MRI (fMRI) studies. The method requires the computation of subject-to-group similarity and within-subject reliability of the observed activations. The idea is that the subject-to-group similarity can inform us about how different each person’s activation map is from the group. However, this information has to be interpreted in the context of the noisiness of each individual map, which can be quantified by assessing its within-subject reliability. Critically, these values can be submitted to a computational model that can assess the relative contribution of group- and subject-level factors to each activation map.

We collected data from a perceptual decision-making task inside an MRI scanner where subjects (N = 50) judged whether a briefly presented display featured more red or blue dots and provided a confidence rating (**Fig. 1A**). The experiment was organized in 96 blocks of 8 trials each (see Materials and Methods for full details). We performed standard analyses to assess the activation maps associated with task trials, as well as with RT and confidence (by comparing trials with higher-vs. lower than the trial-level median RT and confidence). We show that the model can successfully quantify the contribution of group- and subject-level factors to brain activations and that these two factors are approximately equally important in our task.

**Figure 1.**
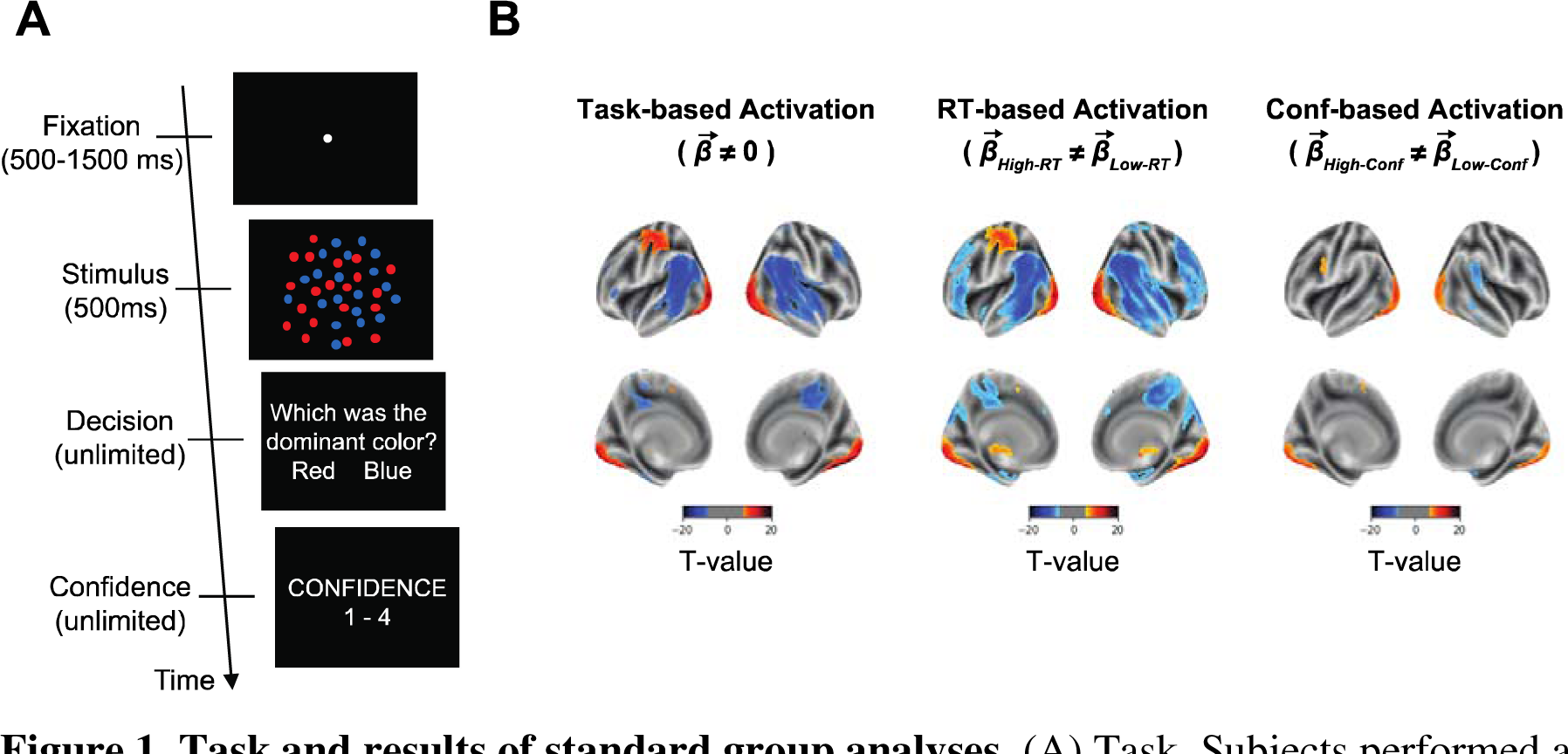
Task and results of standard group analyses. (A) Task. Subjects performed a simple perceptual decision-making task that required them to judge the dominant color in a display of colored dots and rate their confidence. (B) Results of standard second analyses for task-, RT-, and confidence-based contrasts. The analyses showed strong increases and decreases in activation across a range of brain regions for task-(top left), RT- (top middle) and confidence-based (top right) analyses. All maps thresholded at FDR < 0.01 corrected for display purposes.

## Materials and Methods

### Subjects

Fifty-two healthy subjects were recruited for this study. Two subjects were excluded because one had metal braces in their teeth and one decided to stop the experiment after the second run. All analyses were thus based on the remaining 50 subjects (25 females; Mean age = 26; Age range = 19-40; Compensated 20,000 KRW or approximately 18 USD). All subjects were screened for any history of neurological disorders or MRI contraindications. The study was approved by Ulsan National Institute of Science and Technology Review Board (UNISTIRB-20-30-C) and all subjects gave written consent.

### Task

Subjects had to determine which set of colored dots (red vs. blue) was more frequent in a cloud of dots (Fig. 1A). Each trial began with a white fixation dot presented for a variable amount of time between 500-1500 ms at the center of the screen on a black background. Then, the stimulus was shown for 500 ms, followed by untimed decision and confidence screens. The stimulus consisted of between 140 and 190 red- and blue-colored dots (dot size = 5 pixels) dispersed randomly inside an imaginary circle with a radius of 3° from the center of the screen. Four different dot ratios were used – 80/60, 80/70, 100/80, and 100/90, where the two numbers indicate the number of dots from each color. The experiment was organized in blocks of 8 trials each, with each dot ratio presented twice in a random order within a block. The more frequent color was pseudo randomized so that there were equal number of trials where red and blue were the correct answer within a run (consisting of 16 blocks). Subjects used an MRI-compatible button box with their right hand to indicate their decision and confidence responses. For the decision response, the index finger was used to indicate a “red” response and the middle finger for a “blue” response. Confidence was given on a 4-point scale, where 1 is the lowest and 4 is the highest, with the rating of 1 mapped to the index finger and the rating of 4 mapped to the little finger.

Subjects performed 6 runs each consisting of 16 blocks of 8 trials (for a total of 768 trials per subject). Three subjects completed only half of the 6^th^ run and another three subjects completed only the first 5 runs due to time constraints. The remaining 44 subjects completed the full 6 runs. Subjects were given 5 seconds of rest between blocks, and self-paced breaks between runs.

### MRI recording

The MRI data was collected on a 64-channel head coil 3T MRI system (Magnetom Prisma; Siemens). Whole-brain functional data were acquired using a T2*-weighted multi-band accelerated imaging (FoV = 200 mm; TR = 2000 ms; TE = 35 ms; multiband acceleration factor = 3; in-plane acceleration factor = 2; 72 interleaved slices; flip angle = 90°; voxel size = 2.0 x 2.0 x 2.0 mm^3^). High-resolution anatomical MP-RAGE data were acquired using T1-weighted imaging (FoV = 256 mm; TR = 2300 ms; TE = 2.28 ms; 192 slices; flip angle = 8°; voxel size = 1.0 x 1.0 x 1.0 mm^3^).

### MRI preprocessing and general linear model fitting

MRI data were preprocessed with SPM12 (Wellcome Department of Imaging Neuroscience, London, UK). We first converted the images from DICOM to NIFTI and removed the first three volumes to allow for scanner equilibration. We then preprocessed with the following steps: de-spiking, slice-timing correction, realignment, segmentation, coregistration, normalization, and spatial smoothing with 10 mm full width half maximum (FWHM) Gaussian kernel. In control analyses, we used 5 and 20 mm FWHM smoothing to investigate whether the results are due to fine-grained differences in the activations maps between subjects, given that local differences would be substantially reduced by larger smoothing kernels. Despiking was done using the 3dDespike function in AFNI. The preprocessing of the T1-weighted structural images involved skull-removal, normalization into MNI anatomical standard space, and segmentation into gray matter, white matter, and cerebral spinal fluid, soft tissues, and air and background.

We fit a general linear model (GLM) that allowed us to estimate the beta values for each voxel in the brain. The model consisted of separate boxcar regressors for trials that had greater or smaller than the median RT or confidence (trial onset was set to the beginning of fixation and trial offset was set to the confidence response), inter-block rest periods, as well as linear and squared regressors for six head movement (three translation and three rotation), five tissue-related regressors (gray matter, white matter, cerebrospinal fluid, soft tissues, and air and background), and a constant term per run.

### Standard group-level analyses

We first performed a standard group analysis by conducting t-tests across all subjects for each voxel. A task-based analysis compared the obtained beta values with zero to identify regions of activation and de-activation. Two behavior-based analyses compared the beta values for trials with faster-vs. slower-than-median average reaction times (RT) and higher-vs. lower-than-median average confidence. Significance was assessed using p < 0.05 after Bonferroni correction for multiple comparisons. For display purposes, Fig. 1 and Fig. S1 used the more liberal threshold of p < 0.001 uncorrected.

### Within-subject reliability analyses

We examined the within-subject reliability of the whole-brain maps produced by the task, RT, and confidence analyses. To do so, we first re-did each analysis by only using the odd trials, as well as by only using the even trials. We then compared the similarity between the maps obtained for odd and even trials using Pearson correlation. We performed the analysis five times based on the top 10, 25, 50, 75, or 100% of most strongly activated voxels in the following way. We first identified the X% most strongly activated voxels (i.e., the voxels with highest absolute activation values) when only examining the data from the odd trials. The activation values used were the average beta value for task-based analyses, and the t-value (obtained by using a t-test to compare the beta values for trials with above-vs. below-median RT or confidence) for the RT and confidence analyses. This selection procedure ensured that both positively and negatively activated voxels were selected and that an equal number of voxels were selected each time. The activations in the selected top X% of voxels from the odd trials were then correlated with the activations in the same voxels in the even trials, thus obtaining an “odd-to-even” correlation value. Then, using an equivalent procedure, we identified the top X% of most activated voxels in the even trials, and correlated their activations with the activations in the corresponding voxels in the odd trials, thus obtaining an “even-to-odd” correlation value. Finally, we computed the overall within-subject reliability as the average of the odd-to-even and even-to-odd correlation values.

We limited our analysis to a single session because the objective was to develop a method that estimate the contribution of subject- and group-level factors in brain activation using reliability and similarity values. The framework developed here can be extended to include data from multiple sessions but the benefit using a single session is that it will maximize within-session reliability since the reliability between sessions could be affected by multiple exogenous factors (Poldrack et al. 2015; Nakuci et al. 2023).

### Subject-to-group similarity analyses

Critically, we examined the subject-to-group similarity in the maps produced by each analysis. For each subject, we correlated their individual task-, RT-, and confidence-based activation maps with the corresponding group map obtained by averaging the maps of the remaining 49 subjects. Similar to the within-subject reliability analyses, we conducted these analyses separately for the top 10, 25, 50, 75, or 100% of most activated voxels. These voxels were selected in the same way as for the within-subject reliability analyses using all of the data in a given subject; the activations in the voxels identified for a given subject were then correlated with the average activations in the same voxels for the remaining 49 subjects.

### Consistency in activation analysis

As another test of the across-subject similarity, we computed the consistency in the sign of activation. Our main analyses relied on taking correlations, but it is possible that just considering the sign of activation (rather than the strength of activation) would produce different results. To investigate this possibility, we examined the consistency of the sign of voxel activations (positive or negative) across subjects. To do so, we first set all voxels values that were equal to zero to not-a-number value (NaN). This applied to regions that are outside the brain. We then binarized the voxel activation values activation_i_ such that:

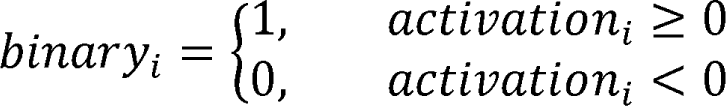

The consistency of the sign of a voxel’s activation across subjects (c_i_) was then calculated as percentage of subjects for which a voxel i was positively or negatively activated using the formula:

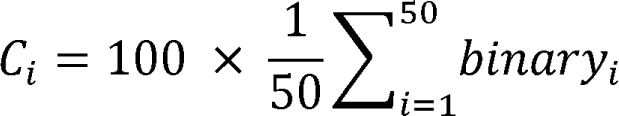

As defined, c_i_ goes from 0 (all subjects having negative activation for that voxel) to 100 (all subjects having positive activations for that voxel), with a value of 50 indicating that half of the subjects had positive and half had negative activation. However, when reporting the values of c_i_, we flipped values under 50 using the formula c_i,flipped_ ; 100 -c_i_, so that these values represent the percent of subjects with negative activations.

The analysis was performed separately for task-based activation maps, RT-based activation maps, and confidence-based activation maps. The activation values were the average beta value (for task-based analyses) or t-value (for RT and confidence analyses).

Low across-subject similarity in these analyses would result in most voxels having consistency, c_i_, values close to 50 (corresponding to the voxel activation having positive sign in half the subjects and negative sign in the other half). However, due to chance, the consistency values are bound to sometimes be higher. Therefore, to enable the appropriate interpretation of the obtained results, we computed the expected consistency values in the maps of 50 subjects whose maps have no relationship to each other.Specifically, we generated a random set of voxel activation values for each of 50 sample subjects. Maximal consistency from the random data was calculated in the same manner as the empirical values and the procedure was repeated 1000 times. This analysis revealed that completely random data would produce a maximal consistency of 80% (for both positive and negative activations) given the number of voxels and number of subjects that we had, which was close to the empirically observed values for RT and confidence analyses.

### Distribution of top-10% most strongly activated voxels

As a final test of the across-subject similarity for the different maps, we sought to identify the consistency of the location of the most strongly activated brain regions across subjects. For each subject, we selected the top-10% most strongly activated voxels by considering the absolute value of either the average beta value (for task-based analyses) or t-value (for RT and confidence analyses). Note that this procedure selected positive and negative activations. We then estimated, for each voxel, the percent of subjects for which the voxel was selected as one of the top-10% most strongly activated voxels. As before, the analysis was performed separately for task-based activation maps, RT-based activation maps, and confidence-based activation maps.

Low across-subject similarity in these analyses would result in most voxels being selected about 10% of the time. However, due to chance, some voxels are bound to be selected more than 10% of the time. Therefore, to enable the appropriate interpretation of the obtained results, we computed the expected level of maximal overlap in the maps of 50 subjects whose maps have no relationship to each other. Specifically, for each of the 50 subjects, we generated a random set voxel activation values. We then selected the top-10% of the highest absolute values from each subject and calculated the overlap across subjects. The expected value from random data was computed as the average maximal overlap after 1000 iterations. This analysis revealed that completely random data would produce a maximal overlap of 28% given the number of voxels and number of subjects that we had, which was only a little less than the empirically observed values for RT and confidence analyses (32% for RT-based analyses and 30% for confidence-based analyses).

### Model specification

The model jointly generates behavior and brain activity maps using minimal assumptions in a way that makes it generalizable across different contexts. The model assumes that the activation map for each trial is a function of seven different factors. The first three are group-level factors (i.e., factors common to all subjects) for the task itself, the influence of RT, and the influence of confidence. The next three factors are subject-level factors (i.e., factors specific to each subject) for the task itself, the influence of RT, and the influence of confidence. Finally, the 7^th^ factor is simply Gaussian noise. Critically, each factor is weighed by a corresponding factor weight that determines the strength of influence of that factor to the final voxel activation values, such that the activation strength (µ) for a given voxel on a given trial is:

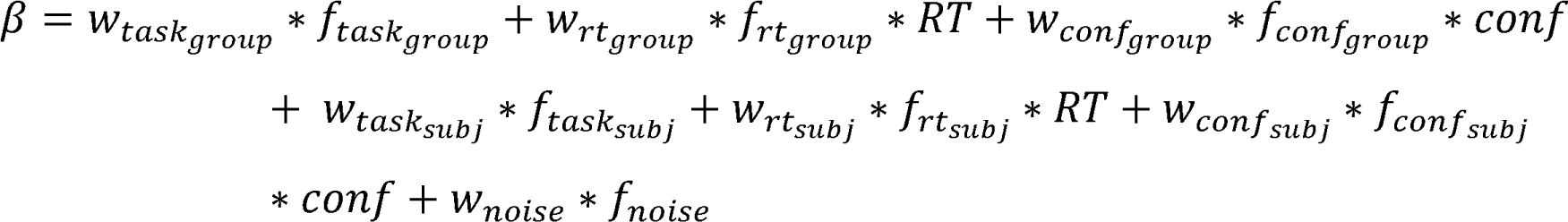

where Kl’ and con1 are the empirical the reaction time and confidence trial, the w’s are the weights associated with each factor, and the 1’s are the factors that influence the voxel activity for a given trial. Without loss of generality, the weight of the noise factor (w_noise_) was fixed to 1. The 1 variables are the component of activation that influence the voxel activity for a given trial and *f* can be thought of as the latent (unobserved) component of activation that is associated with the task, RT, confidence, and noise. The value of each factor 1 was randomly sampled from a standard normal distribution such that group-level factors were randomly sampled for each voxel, subject-level factors were randomly sampled for each voxel and subject, and the noise factor was randomly sampled for each voxel, subject, and trial. We note that the model does not predict beta values for individual regressors. Instead, it generates beta values that already take into account all regressors, which are then used to compute subject-to-group similarity and within-subject reliability values.

The advantage of a model-based approach is that (1) it provides the ratio of subject to group level contribution and (2) it allows us to compare the contribution of subject-and group-level factors relative to the noise in the data. Alternatively, the ratio can be calculated directly from the within-subject reliability and subject-to-group similarity, but the advantage of the model is that it allows us to compare the group- and subject-level factors to the noise level. Therefore, a model-based approach allows for a more thorough analysis of the contribution of subject- and group-level factors to the brain activation.

### Model fitting

We first fit the model to the empirically observed within-subject reliability and subject-to-group similarity values. The model had six free parameters corresponding to the weights, w, of the group- and subject-level factors that determined the simulated µ value for each voxel in each trial. For a given set of weights, we simulated a complete experimental dataset by generating simulated data for 50 subjects with 768 trials per subjects. Based on these data, we then computed the within-subject reliability and subject-to-group similarity values in the same way as for the empirical data. When simulating the model, we observed that the exact number of voxels used made no systematic difference to the observed values of the obtained within-subject reliability and subject-to-group similarity values. Therefore, we used 10,000 voxels, which allowed for stable values to be obtained on different iterations. The fitting minimized the mean squared error (MSE) between the simulated and empirically observed within-subject reliability and subject-to-group similarity values calculated using the top-100% most activated voxels (that is, using all voxels). Once the fitting was completed, we also generated the predictions of the best-fitting model for the within-subject reliability and subject-to-group similarity values calculated using the top 10, 25, 50, and 75% most activated voxels. The fitting itself was carried out using the Bayesian Adaptive Direct Search (BADS) toolbox (Acerbi and Ma 2017). We fit the model 10 times are reported the best fitting model among the 10 iterations. We repeated the model fitting 10x to avoid local minima when estimating parameters, as is standard practice in the field and our laboratory (Shekhar and Rahnev 2020; Yeon and Rahnev 2020).

### Model Comparison

We have compared the Full model (Subject + Group + Noise factors) with a Subject-Only model (Subject + Noise factors) and Group-Only model (Group + Noise factors). We simulated each model 25x and calculated the mean-squared error (MSE) between the model-based and empirical values for the subject-to-group similarity and within-subject reliability values. In addition, we compared the different models using Akaike Information Criterion (AIC) and Bayesian Information Criterion (BIC).

### Data and code availability

Processed data and code are available at https://osf.io/gyw8f/.

## Results

We first performed standard group fMRI analyses by conducting t-tests across all subjects for each voxel. We found that contrasts related to the task (Task > Background), RT (Fast RT > Slow RT), and confidence (High confidence > Low confidence) all produced regions of strong activation and de-activation (Fig. 1B). We inspected the activations for task, RT, and confidence in subjects 1-3 and found that all three subjects demonstrated relatively consistent activation patterns (Fig. 2). However, there appeared to be consistent across-subject differences in the activation maps, which could not be attributed purely to noise as they also appeared in maps produced by only the odd or only the event trials for a given subject. These results hint at the idea that both group- and subject-level factors may be contributing to the observed activations.

**Figure 2.**
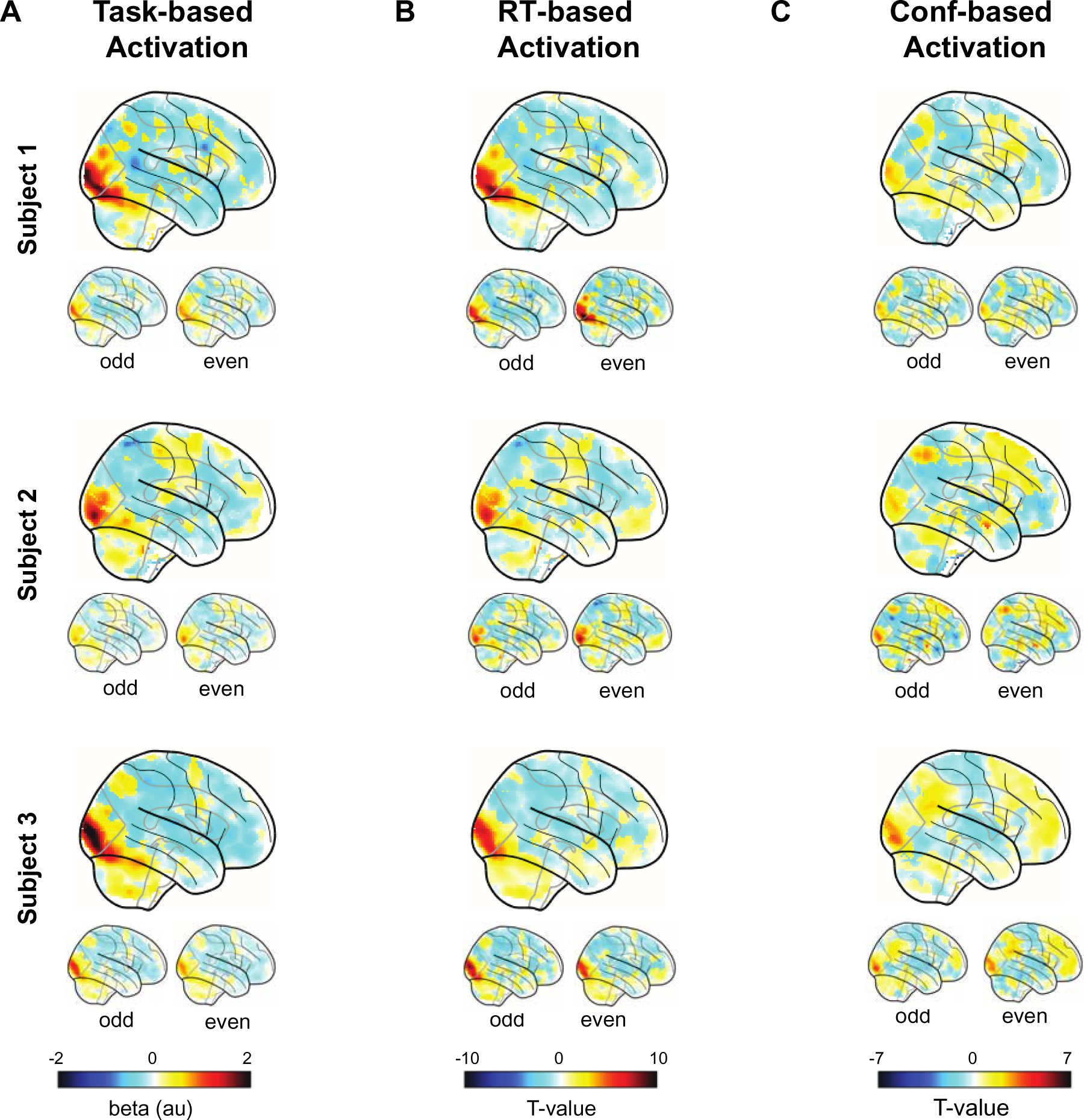
Trial-level activations for task, RT, and confidence for three example subjects. Trial-level activation maps for (A) task, (B) RT, and (C) confidence contrasts from the first three subjects. Small brains underneath represent the same contrasts conducted only on odd or even trials. Similar activations for all three subjects appear for all trial-level contrasts.

To formally test these impressions, we first examined both the within-subject reliability and subject-to-group similarity of the whole-brain maps for the task, RT, and confidence contrasts. We computed within-subject reliability by performing Pearson correlations between the activations obtained when examining only the odd or only the even trials. We computed subject-to-group similarity by correlating each subject’s brain map with the group map obtained by averaging the maps of the remaining 49 subjects.

As may be expected from Figure 2, for task-based activation we found strong within-subject reliability (r_act_ = 0.81 ± 0.013, p < 0.001) and subject-to-group similarity in task activations (r_act_ = 0.72 ± 0.013, p < 0.001; Fig. 3A). In the same manner, RT- and confidence-based maps exhibit strong within-subject reliability (r_rt_ = 0.74 ± 0.014, p < 0.001; r_conf_ = 0.55 ± 0.028, p < 0.001; Fig. 3B-C, top). Critically, we examined the subject-to-group similarity for the RT and confidence maps. Echoing the qualitative impressions from Figure 2, we found a high degree of similarity across subjects for the RT-based maps (r_rt_ = 0.69 ± 0.014, p < 0.001; Fig. 3B, **bottom**) and confidence-based maps (r_conf_ = 0.52 ± 0.025; Fig. 3C, **bottom**).

**Figure 3.**
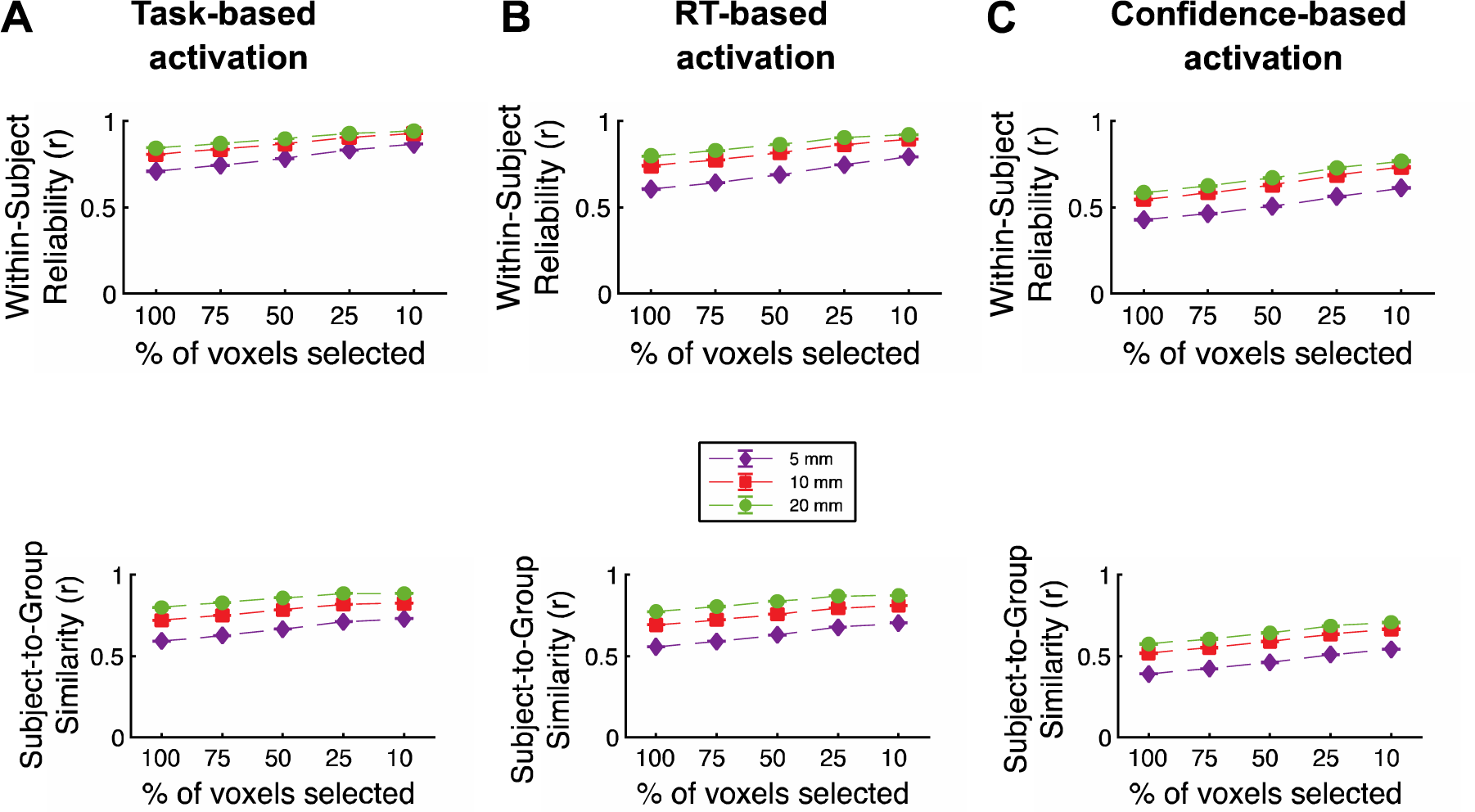
Within-subject reliability and subject-to-group similarity. Within-subject reliability and subject-to-group similarity values as a function of the percent of most activated voxels selected for (A) task- (B) RT- and (C) confidence-based activation.

Subject-to-group similarity is computed as the average similarity between the maps of each person and the group map of the remaining subject. Error bars show SEM.

One potential concern with these types of analyses could be that they may be biased by voxels that are either particularly noisy or not involved in the task in any way. Therefore, to test whether these results are robust, we repeated them by first selecting the top 75, 50, 25, or 10% of the most strongly activated voxels for each subject (see Methods). These analyses showed that selecting smaller percentages of the most highly activated voxels generally increased both the within-subject reliability and subject-to-group similarity, but the pattern of results remained essentially unchanged.

To gain further intuition for the underlying effects, we conducted two additional analyses. First, we tested the consistency of the sign of voxel activations (whether they were positive or negative) across subjects. We found that for the task maps, there were large portions of the brain that showed consistently positive or consistently negative activations (Fig. 4A **left**). Indeed, the maximal overlap across subjects was 100% for both positive and negative activations. Further, we found many areas of strong consistency with maximal overlap of 100% and 98% for positive and negative activations in the RT maps, and maximal overlap of 100% and 88% for positive and negative activations in the confidence maps (Fig. 4A **middle and right**). (Note that the expected values in random data are 80% for positive and negative activations)

**Figure 4.**
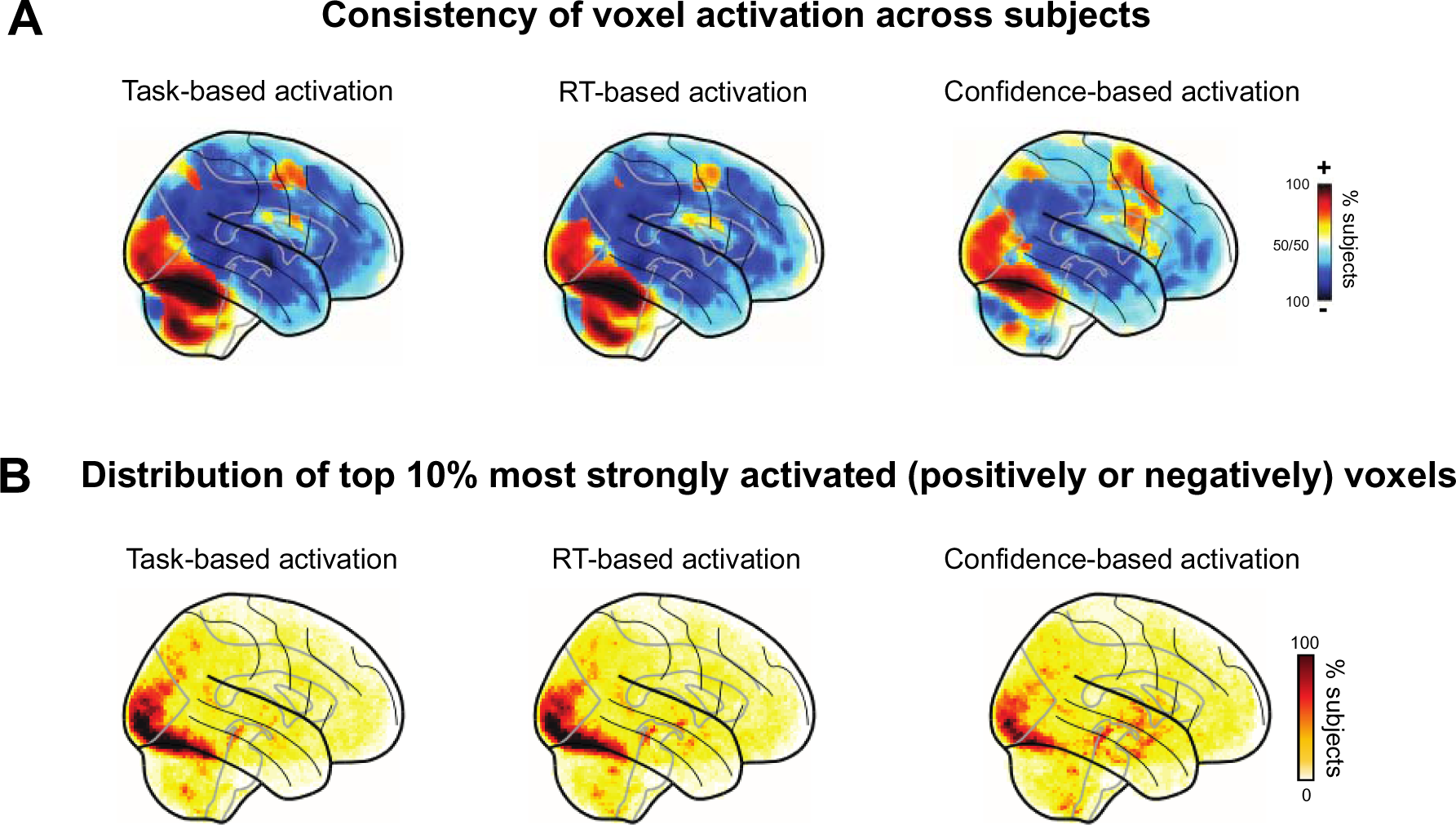
Maps of the activation consistency and distribution of the top-10% most activated voxels across subjects. (A) Maps of voxel consistency computed as the proportion of subjects showing a positive or negative relationship between voxel activity and behavior. Task activations maps, as well as RT and confidence maps show a high level of consistency. (B) Maps of the distribution of the top-10% most activated voxels. Task activations maps, as well as RT and confidence maps contain areas with a high level of consistency in occipital and parietal lobes.

Second, we examined the distribution of the locations of the top-10% most strongly activated voxels for each subject (both positive and negative activations were considered). The most strongly activated voxels clustered in the occipital and parietal lobes (Fig. 4B **left**). The maximum overlap among the 10% most activated voxels across subjects was 98%. Further, there were again areas of strong clustering of the most activated voxels (mostly in the occipital lobe) for both RT and confidence maps (maximal overlap: 98% and 78%, respectively; Fig. 4B **middle and right**). Altogether, both additional analyses further underscore the high level of consistency for task, RT and confidence maps across subjects. We also repeated the same analyses above with a wide range of smoothing levels (from 5 to 20 mm) and obtained very similar results (**Fig. S1 and S2**).

Having quantified the within-subject reliability and subject-to-group similarity between different types of analyses, we used this information to quantify the contribution of group- and subject-level factor by building a simple computational model. The critical idea behind the model is to separately model group-level factors (i.e., factors that are identical for all subjects) and subject-level factors (i.e., factors that are different for each subject). The inputs into the model are the empirical within-subject reliability and subject-to-group similarity values, as well as the empirical RT and confidence values.

The simulation generates idealized beta values (voxel activations) for each trial characterized by a given RT and confidence values. Note that the activations produced by the model are not mapped onto specific voxels in the brain and do not form a meaningful spatial map. That is, to keep the model simple, individual voxel activation for each group- and task-level factor were generated randomly by ignoring known temporal and inter-regional dependencies.

Critically, the model produces the idealized beta values based on three group-level factors (group task map, group RT map, and group confidence map), three subject-level factors (subject-specific task map, subject-specific RT map, and subject-specific confidence map), and one noise factor (Fig. 5A). The weight of the noise factor was fixed to 1, leaving the model with a total of six free parameters (one for the weight of each group- and subject-level factor). We then fit the within-subject reliability and subject-to-group similarity produced by the model to the observed values computed using 100% of the voxels.

**Figure 5.**
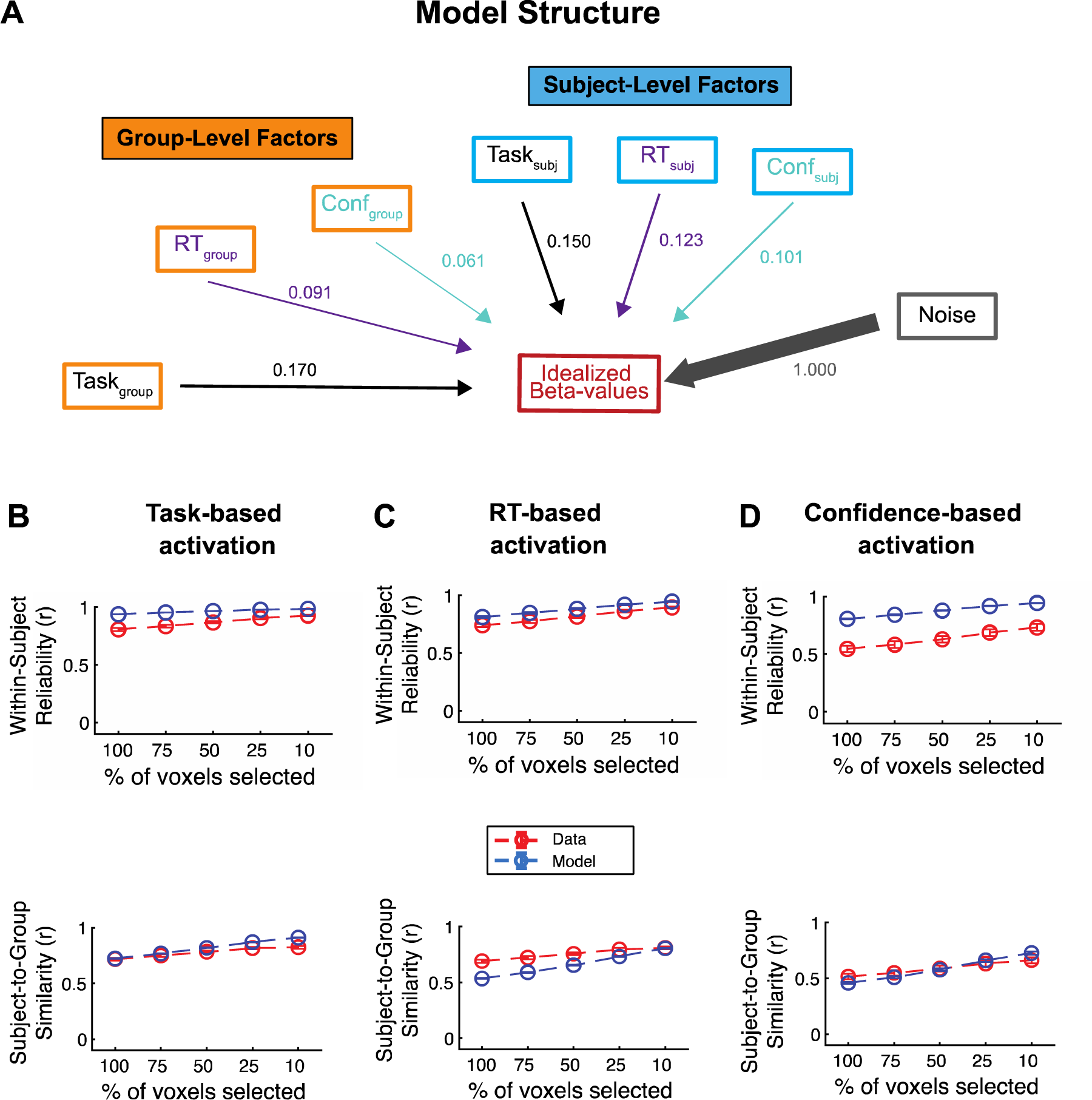
Model structure and model fits. (A) Graphical depiction of the model at the trial level. The model generates an idealized set of beta values for an individual trial as the confluence of three group-level, three subject-level, and one noise factor. The thickness of the arrows and associated numbers correspond to the weights obtained from fitting the model to the data. (B-C) Model fits to the within-subject reliability (*top*) and subject-to-group similarity (*bottom*) values for (B) task, (C) RT, and (D) confidence analyses. The model was fit only to the empirical data with 10-mm smoothing where 100% of voxels selected. Despite its simplicity, the model is able to reproduce the empirical data for the remaining analyses with smaller percentages of selected voxels very well.

Despite its simplicity, the model was able to provide excellent fit to the data from Fig. 3 by capturing closely the observed patterns of within-subject reliability (Fig. 5B**-D**, **top**) and subject-to-group similarity (Fig. 5B**-C**, **bottom**) for the data with 10 mm smoothing. We also separately fit the data with 5 and 20 mm smoothing and obtained equally good fits.

Critically, the model allowed us to examine the weights of the group- and subject-level factors, thus providing insight into the relative contribution of each. We found slightly larger contribution weights for the group-than subject-level task factors (subject-level factor weight = 0.150, group-level factor weight = 0.170, ratio = 0.88; Fig. 6A, B). Thus, the group-level factors were only slightly higher than the subject-level factors, pointing to a balance between influences that are common across all subjects and influences that are specific to each individual. On the other hand, we observed slightly higher relative weights for the subject-level factors for the RT and confidence maps at the trial level (RT: subject-level factor weight = 0.123, group-level factor weight = 0.091, ratio = 1.35; Confidence: subject-level factor weight = 0.101, group-level factor weight = 0.061, ratio = 1.65). In other words, our model suggests that group- and subject-level factors have relatively similar influence on task activation maps, which corresponds well to recent findings about group- and subject-level influences on brain connectivity (Gratton et al. 2018). However, the relative contribution of all group- and subject-level factors is small relative to the contribution of noise (Fig. 6C).

**Figure 6.**
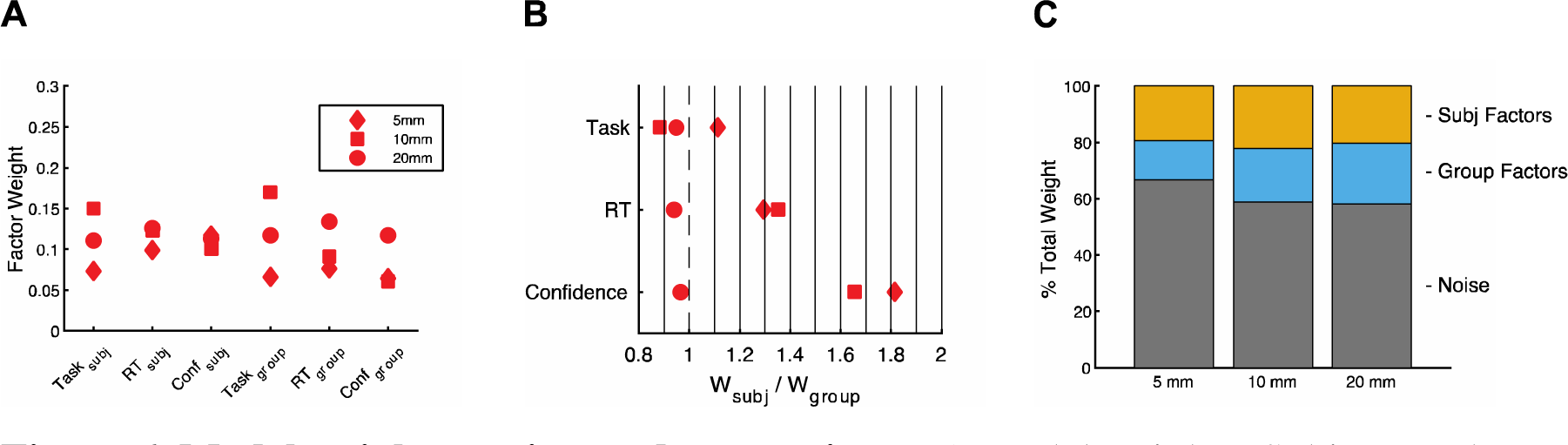
Model weights, ratios, and proportions. A) Model weights. Subject- and group-level weights obtained from fitting the model separately to each level of smoothing (5, 10, and 20 mm). B) Weight ratios. Relative weights of the subject-level and corresponding group-level factors from each analysis. C) Factor proportions. The combined percent accounted for by subject, group, and noise factors contributing to the activation on an individual trial. Subject and group factors reflect the summed task, RT, and confidence weights.

To examine the robustness of the modeling results, we repeated the model fitting on data with 5 to 20 mm smoothing. These two additional analyses produced similar results: the weights ratio between the subject- and group-level factors was between 0.8 and 1.21 for the task factors in all cases, between 0.9 and 1.4 for RT, and between 0.9 and 1.8 for confidence (Fig. 6B).

Additionally, we compared the Full model (Subject + Group + Noise factors) with a Subject-Only model (Subject + Noise factors) and a Group-Only model (Group + Noise factors) (**Fig. S3A-C**). We simulated each model 25x and calculated the mean-squared error (MSE), Akaike Information Criterion (AIC), and Bayesian Information Criterion (BIC) between the model-based and empirical within-subject reliability and subject-to-group similarity values. The reliability and similarity values estimated from the Full model exhibited lower MSE, AIC and BIC values compared to the Subject-Only or Group-Only models (paired sample t-test, p < 10^-26^; **Fig. S3D-F**). These results indicate that there are both subject and group components in both task-and behavior-based brain activation maps.

Lastly, we explored whether we would obtain similar results if we repeat these analyses at the level of blocks (of eight trials each) rather than trials. Similar to the trial-level analyses, we found relatively high subject-to-group similarity and within-subject reliability values for task activations. However, analyses of average RT and confidence on the block level revealed very low subject-to-group similarity values but reasonably high within-subject reliabilities, which was reflected in much higher values for subject-compared to group-level factors in our model (**Fig. S4-S8**). These results suggest that other types of analyses than the standard ones included here may result in different contributions of subject- and group-level factors.

## Discussion

A major goal of neuroscience research has been to understand the neural correlates of behavior. Behavior is a complex phenomenon that is often specific to a person (Eilam 2015; Forkosh et al. 2019). Idiosyncratic behavioral responses are ubiquitous in social situations (Durlauf 2001), economic decisions (Kable and Glimcher 2007), judgments of beauty (Martinez et al. 2020), confidence ratings (Navajas et al. 2017), response bias (Rahnev 2021), and even low-level perception (Afraz et al. 2010). Here we develop a method to quantify the level of idiosyncrasy in brain activations by estimating the relative contributions of group- and subject-level factors. By applying this method to a new dataset where subjects (N=50) completed a perceptual decision-making task, we find that for standard analyses at the trial level, the influence of subject-level factors is only slightly stronger than the influence of group-level factors.

There are at least two important conclusions that one can draw from the current results. First, across all analyses performed here, subject-level factors were at least as important as group-level factors. While this effect could be at least partly driven by issues such as misalignment across different brains, the results were remarkably stable whether they were computed using 5-, 10-, or 20-mm smoothing. If brain misalignment were the main source of the observed idiosyncrasy here, one would expect that larger smoothing would produce different results. These results suggest that idiosyncratic, subject-level factors may play a large role in observed brain activations. Our findings thus highlight the need for a renewed focus on investigating the brain-behavior relationship at the level of single subjects (Gilmore et al. 2021; Gordon and Nelson 2021; Naselaris et al. 2021; Song and Rosenberg 2021).

Our current results also suggest novel ways for finding robust biomarkers for various mental disorders (Kaufmann et al. 2017; Elliott et al. 2018; Li et al. 2020; Parkes et al. 2020). Most research in the field has focused on biomarkers unrelated to behavior such as functional connectivity patterns at rest (Woodward and Cascio 2015; Drysdale et al. 2017). An exciting possibility is that subject-level activations maps for disease-relevant behaviors could serve as much more powerful biomarkers because of their high reliability and clear differences among people. Focusing directly on the relationship between one’s behavior and one’s brain activations may help to delineate the intricate relationship between the brain and psychopathology (Gratton et al. 2020). Therefore, subject-level effects could be crucial to diagnosing and treating different mental illnesses. Additionally, an analysis that is focused on subject-level variability might be more informative since between-subject analyses ignore the large degree of within-subject variability (Fisher et al. 2018; Lebreton et al. 2019).

It is worth noting that contribution of group- and subject-level factors might change. In some tasks, the group-level factors might play a larger role, whereas in other tasks the subject-level factor might play a larger role. These different tasks might be valuable for isolating the group- and subject-level components of cognitions. Future research should estimate the contribution of these factors in a wider variety of tasks and contrasts.

Previous work has utilized mixed-effect modeling to estimate the contribution of subject- and group-factors (Woolrich et al. 2004; Friston et al. 2005; Chen et al. 2013). This prior work has relied on estimating these effects directly from the underlying brain activation patterns associated with given condition. The framework developed here builds upon this work to simulate brain activation to estimate the contribution of subject- and group-level factors. In a similar fashion to previous work, the subject-level factors can be thought of as random effects and the group-level factors as fixed effects.

Despite the fact that our model is able to fit the data quite well, it is nonetheless important to highlight the model’s limitations. In particular more complex models such as hierarchical models might perform better. However, we are not able to fit a more complex model because we are fitting group-level data (e.g., average subject-to-group similarity values) rather than each individual separately. A second limitation pertains to whether the observed subject-level effects are stable across multiple sessions. In the current analysis, we used fMRI data from a single session, but fMRI signals are highly variable between sessions even for the same subject (McGonigle et al. 2000; Zandbelt et al. 2008). Future studies should utilize multiple sessions to confirm the stability of the subject-level effects. Third, nearby voxels are known to be related to each other, thus resulting in substantial spatial autocorrelations in fMRI (Shinn et al. 2023). Our analyses do not account for such spatial autocorrelations because they do not attempt to generate voxel-level predictions. Nonetheless, it could be useful for future models to include such autocorrelations. Fourth, in our analyses we split trials based on the median, but median split can have undesirable statistical properties. An alternative would be to use parametric modulation to estimate the relationship between brain activation and RT and confidence.

In conclusion, we develop a computational model to quantify the contribution of group- and subject-level factors in activation patterns. Our model suggests that activations related to task, RT, and confidence in a perceptual decision-making task are influenced equally strongly by group- and subject-level factors. However, both group- and subject-level factors are dwarfed by a noise factor. Taken together, our method provides a more detailed understanding of the idiosyncrasy levels in brain activations.

## Acknowledgments

This work was supported by the National Institutes of Health (grant R01MH119189 to DR) and the Office of Naval Research (grant N00014-20-1-2622 to DR).

## Author contributions

Conceptualization: JN, DR; Methodology: JN, JY, KX, DR; Data Curation: JY, JHK, SPK; Visualization: JN, DR; Funding acquisition: DR; Writing – original draft: JN, DR; Writing – review & editing: JN, JY, KX, JHK, SPK, DR

---

**Fig. S1.**
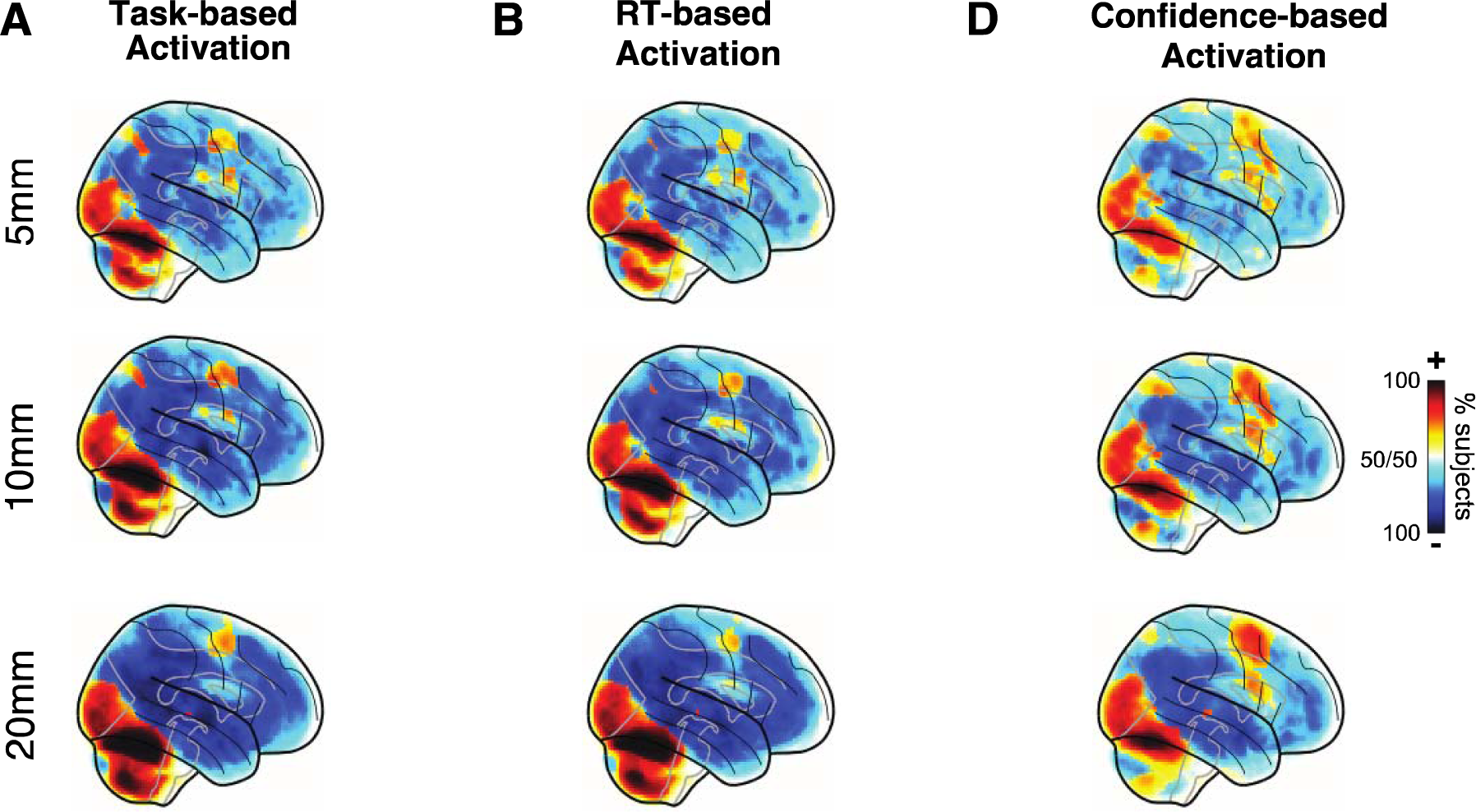
Trial-level analysis maps of voxel activation consistency across subjects. A) Task-based activation. B) RT-based activation. C) Confidence-based activation. All maps exhibited strong areas of consistency. Analysis was conducted on fMRI data smoothed with 5, 10, and 20 mm FWHM kernels. The 10 mm results are the same as in the main manuscript and are shown here for comparative purposes. Again, similar results are obtained for different levels of smoothing.

**Fig. S2.**
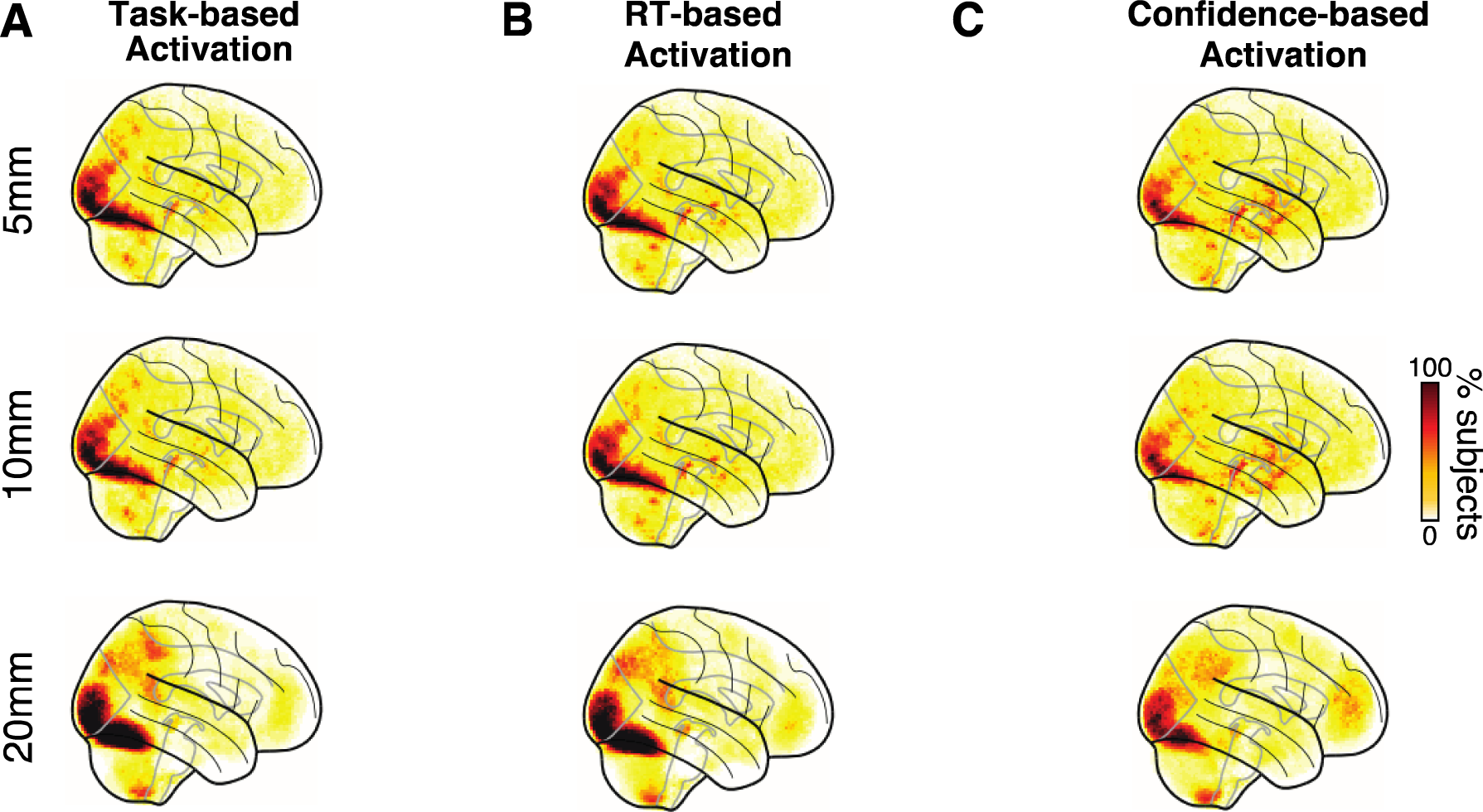
Trial-level maps of the distribution of the top-10% most activated voxels. A) Task-based activation. B) RT-based activation. C) Confidence-based activation. All maps exhibited strong areas of consistency compared. Analysis was conducted on fMRI data smoothed with 5, 10, and 20 mm FWHM kernels. The 10 mm results are the same as in the main manuscript and are shown here for comparative purposes. Again, similar results are obtained for different levels of smoothing.

**Fig. S3.**
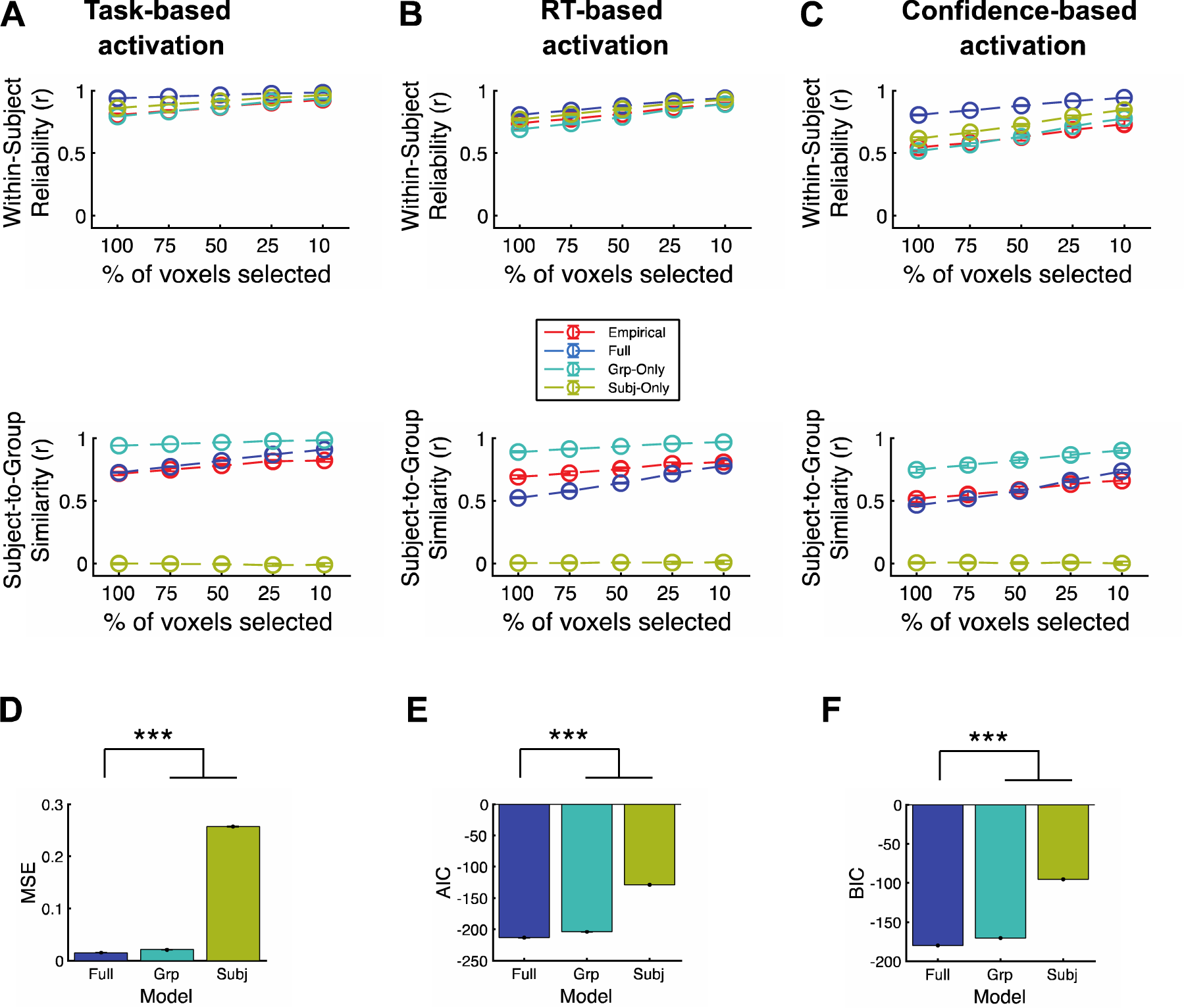
Comparing the Full, Subject-Only, and Group-Only models. Sample within-subject reliability and subject-to-group similarity from the simulation using the Full, Subject-Only, and Group-Only factors in the simulation for (A) task-, (B) RT-, and (C) confidence-based activations. The full simulation model used subject-, group-, and noise-factors. The Subject-Only simulation model used subject and noise factors. The Group-Only simulation model used group and noise factors. (D-F) Model performance. The within-subject reliability and subject-to-group similarity values estimated in 25 simulations, (D) the mean-squared error (MSE), (E) AIC, and (F) BIC were estimated by comparing the within-subject reliability and subject-to-group similarity from the simulation with the empirical values. The Full model outperformed both the Subject-Only and Group-Only models. Error bars show SEM. *** p < 0.001.

**Fig. S4.**
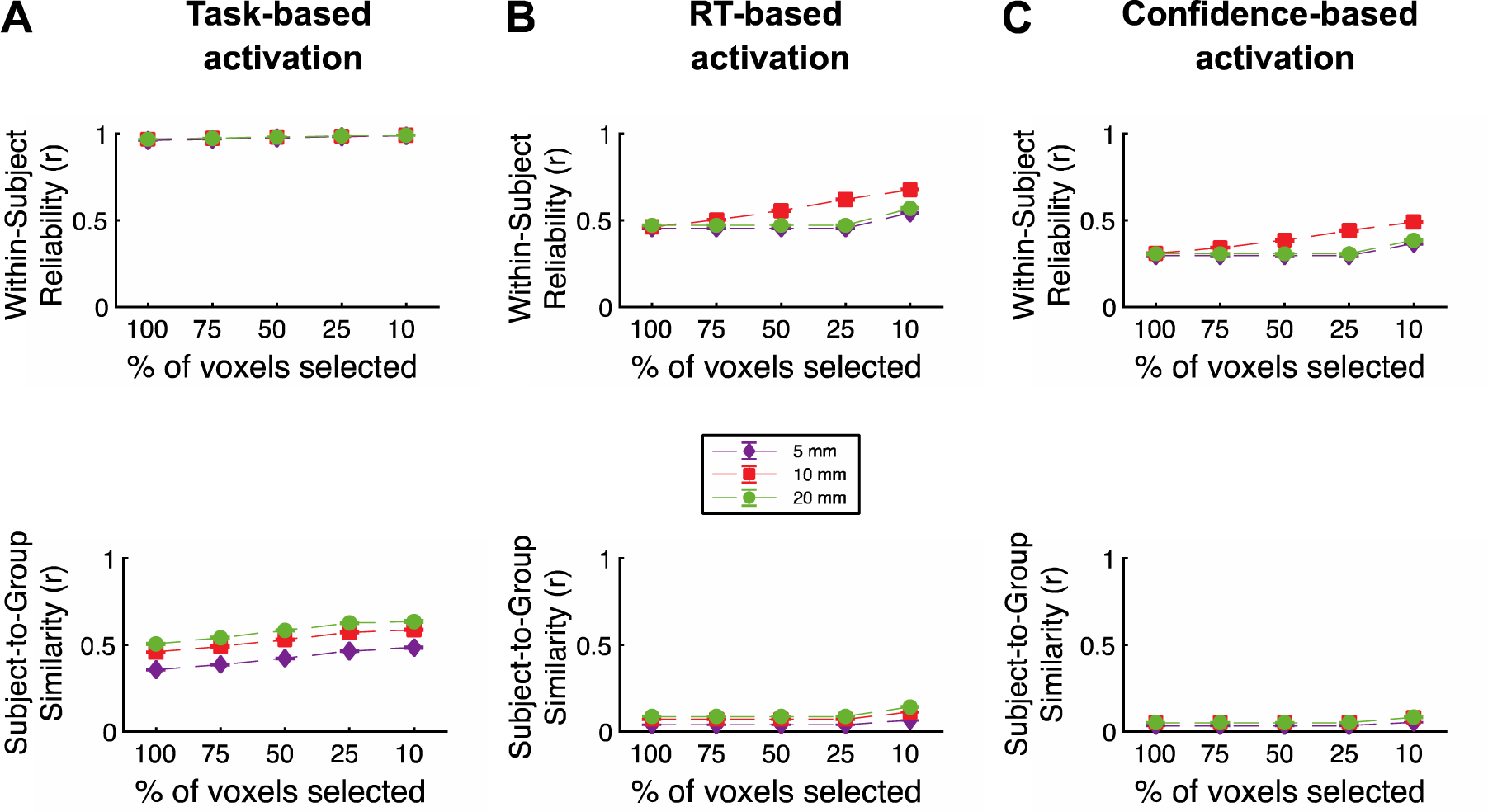
Within-subject reliability and subject-to-group similarity for analyses conducted at the block level. Within-subject reliability and subject-to-group similarity values of the whole-brain maps produced by the (A) task-, (B) RT-, and (C) confidence-based analyses. We fit a general linear model (GLM) that allowed us to estimate the beta values for each voxel in the brain. For the block-analyses, the model consisted of regressors for each individual block (block onset was set to the beginning of fixation on the first trials and block offset was set to the confidence response of the last trial in the block), inter-block rest periods, as well as linear and squared regressors for six head movement (three translation and three rotation), five tissue-related regressors (gray matter, white matter, cerebrospinal fluid, soft tissues, and air and background), and a constant term per run. Two behavior-based analyses compared the beta values for blocks with faster-vs. slower-than-median average reaction times (RT) and higher-vs. lower-than-median average confidence. Within-subject reliability and subject-to-group similarity of the whole-brain maps produced by the task, RT, and confidence analyses was examined in the same manner as for the trial level analysis. The fMRI data were spatially smoothed with 5 mm, 10 mm, or 20 mm full width half maximum (FWHM) Gaussian kernel. As can be observed, very similar results are obtained for different levels of smoothing, indicating that the results obtained are likely due to large-scale rather than small-grained differences in the maps. Error bars show SEM.

**Fig. S5.**
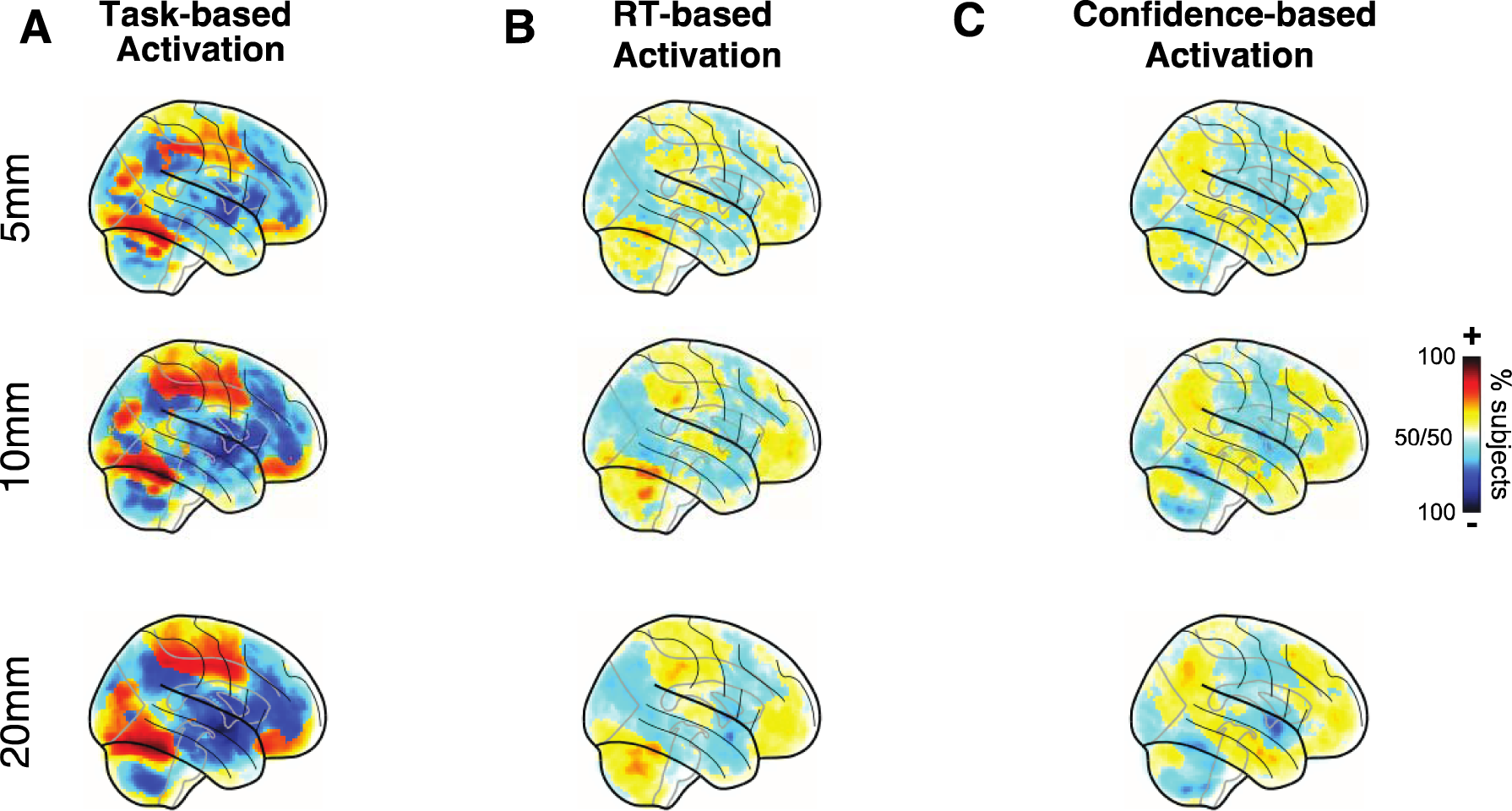
Block-level maps of voxel activation consistency across subjects. A) Task-based activation. B) RT-based activation. C) Confidence-based activation. Task-based activations exhibited strong areas of consistency, but both the RT and confidence maps showed much weaker consistency across subjects. Analysis was conducted on fMRI data smoothed with 5, 10, and 20 mm FWHM kernels. Again, similar results are obtained for different levels of smoothing.

**Fig. S6.**
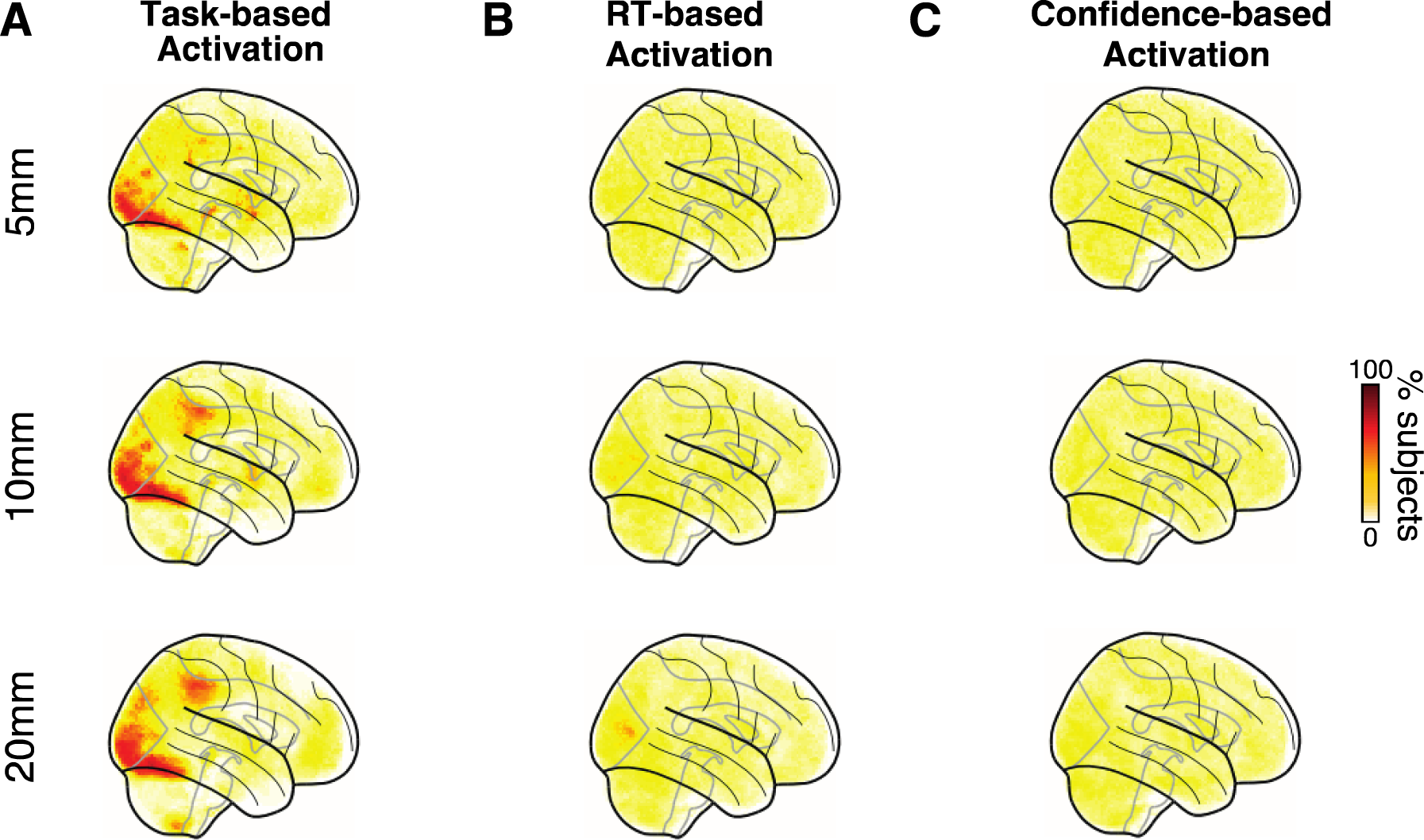
Block-level maps of the distribution of the top-10% most activated voxels. A) Task-based activation. B) RT-based activation. C) Confidence-based activation. Task-based activations exhibited strong areas of consistency, but both the RT and confidence maps showed much weaker consistency across subjects. Analysis was conducted on fMRI data smoothed with 5, 10, and 20 mm FWHM kernels. Again, similar results are obtained for different levels of smoothing.

**Figure S7.**
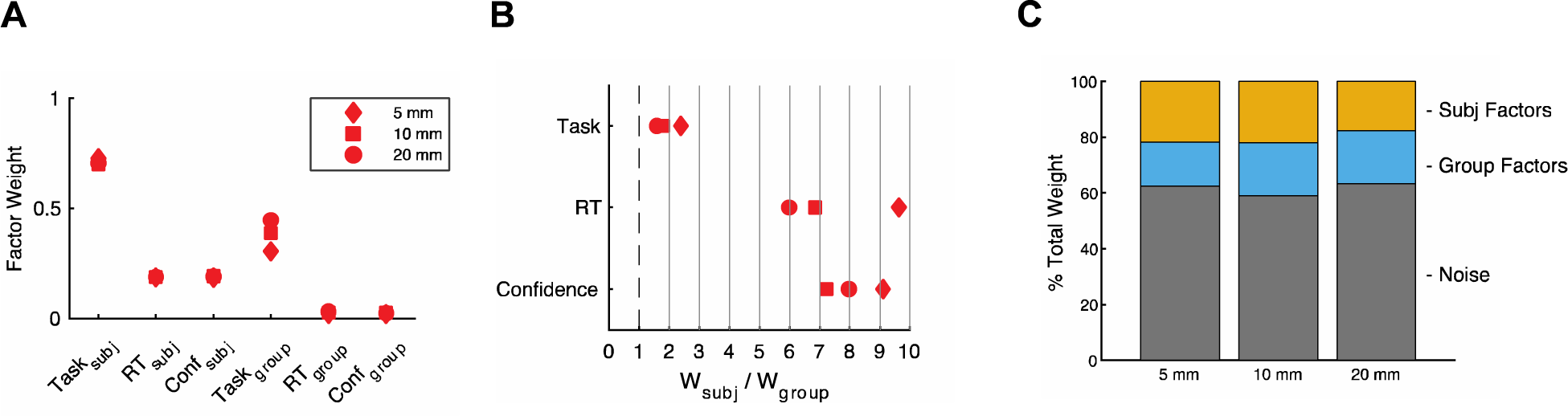
Block-level model weights, ratios, and proportions. A) Model weights. Subject- and group-level weights obtained from fitting the model separately to the data with each smoothing level. B) Weight ratios. Relative weights of the subject-level and corresponding group-level factors for each smoothing level. C) Factor proportions. The relative weight of subject, group, and noise factors contributing to the activation on an individual block. Subject and group factors reflect the summed task, RT, and confidence weights.

**Fig S8.**
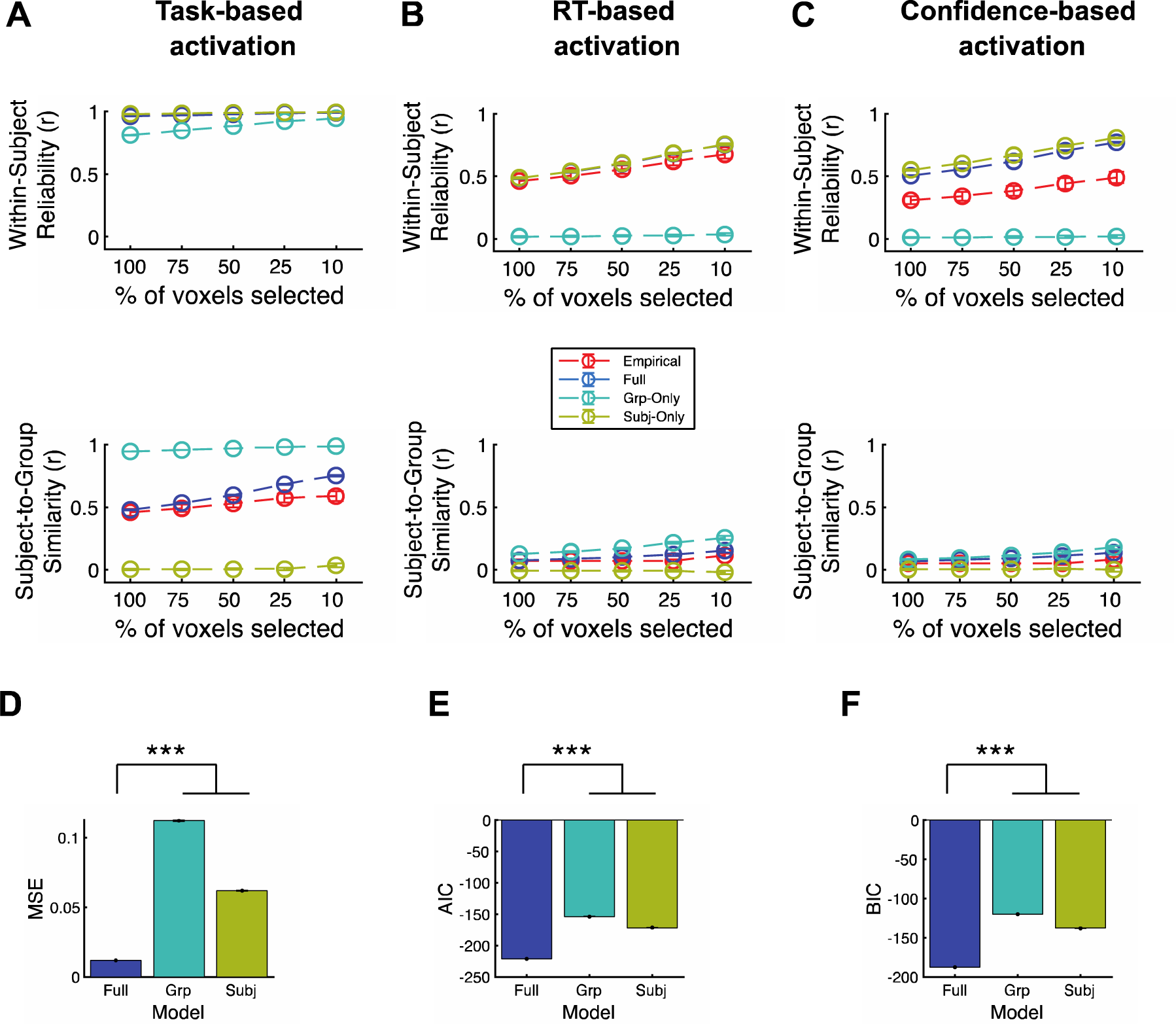
Comparing the Full, Subject-Only, and Group-Only models for block-level analysis. Sample within-subject reliability and subject-to-group similarity from the simulation using the Full, Subject-Only, and Group-Only factors in the simulation for (A) task-, (B) RT-, and (C) confidence-based activations. The full simulation model used subject-, group-, and noise-factors. The Subject-Only simulation model used subject and noise factors. The Group-Only simulation model used group and noise factors. (D-F) Model performance. The within-subject reliability and subject-to-group similarity values estimated in 25 simulations, (D) the mean-squared error (MSE), (E) AIC, and (F) BIC were estimated by comparing the within-subject reliability and subject-to-group similarity from the simulation with the empirical values. The Full model outperformed both the Subject-Only and Group-Only models. Error bars show SEM. *** p < 0.001.

